# Live-cell single-molecule analysis of β_2_-adrenergic receptor diffusion dynamics and confinement

**DOI:** 10.1101/406488

**Authors:** Niko Schwenzer, Hendrik Bussmann, Sebastian Franken, Hanns Häberlein

## Abstract

Signal transduction mechanisms and successive regulatory processes alter the lateral mobility of β_2_-adrenergic receptors (β_2_AR). In this work we combined modern single particle tracking methods in order to analyze the diffusion dynamics of SNAP-tagged β_2_AR in HEK wild-type cells and HEK β-arrestin knockout cells before and after agonist stimulation. For analysis of trajectories we first used mean squared displacement (MSD) analysis. Secondly, we applied an advanced variational Bayesian treatment of hidden Markov models (vbSPT) in combination with the recently introduced packing coefficient (Pc), which together provided a detailed model of three discrete diffusive states, state transitioning and spatial confinement. Interesting to note, state switching between S3 (fast-diffusing) and S1 (slow-diffusing) occurred sequentially over an intermediate state S2. After ligand stimulation more SNAP-tagged β_2_AR in HEK wild-type cells switched occupancy into the slow-diffusing state, whereas less receptors were found in the fast diffusive state. Unexpectedly, all three states showed a fraction of confined receptor mobility that increased under stimulation, but confinement sizes were unaffected. Receptor diffusion characteristics were comparable in HEK β-arrestin knockout cells under basal conditions and only minor but non-significant changes occurred upon stimulation, as expected from the depletion of β-arrestin, an important regulatory protein. The data presented here on the occurrence of different diffusion states, their transitioning and variable spatial confinements clearly indicate that lateral mobility of β_2_AR is much more complex than previously thought.

## Introduction

G protein-coupled receptors (GPCRs) represent one of the largest protein families in the mammalian genome^1^. GPCRs are involved in numerous important signaling processes initiated e.g. by neurotransmitters or hormones. Major mechanisms and regulation of GPCR signal transduction have been discovered^2,3^. One of the most investigated GPCR is the β_2_-adrenergic receptor (β_2_AR). Stimulation of β_2_AR activates a heterotrimeric G protein, from which the G_s_ alpha subunit is released to activate adenylate cyclase (AC). AC catalyses the synthesis of the second messenger cyclic adenosine monophosphate (cAMP). Desensitization of β_2_AR, achieved by two phosphorylation steps mediated by protein kinase A (PKA) and G protein-coupled receptor kinase 2 (GRK2), respectively, prevents over-stimulation of the cell. Furthermore, redistribution of phosphorylated β_2_AR from functional microdomains into clathrin coated pits followed by receptor internalization reduces receptor density on the cell surface.

However, little is known about the lateral mobility of β_2_AR during signal transduction and regulation processes. Immobile, confined, and free diffusion behaviour were found possibly characterizing different functional states of adrenergic receptors^4–6^. Sungkaworn et al. recently observed immobilization of α_2A_-adrenergic receptors while interacting with G_s_ alpha subunits^4^. Interactions of β-adrenergic receptors with scaffold proteins^7^ or confinement in caveolae domains^8^ are further discussed for varying receptor diffusion properties. Furthermore, desensitized β_2_AR interacting with regulatory proteins like β-arrestin, AP-2, and dynamin recruited to clathrin coated pits leads to the formation of an internalization complex^9,10^ with nearly immobile lateral diffusion behavior.

In this work we applied modern single particle tracking methods to analyze the lateral mobility of SNAP-tagged β_2_AR in HEK wild-type cells and HEK β-arrestin knockout cells. The latter cell line lacks expression of regulatory adaptor proteins β-arrestin2 and β-arrestin3^9^. Fluorescent labeling of β_2_AR was realized by expression of SNAP-tag fusion protein^11^ in both cell lines combined with the highly specific and photostable dye substrate BG-CF640R^12,13^. Distributions of single β_2_AR diffusion coefficients derived from linear fitting of MSD curves of each track were determined. Additionally, all tracks of each condition were evaluated using a software based on a variational Bayesian treatment of Hidden Markov Models (vbSPT) to identify discrete diffusive states and state transitioning probabilities. We also implemented the recently introduced packing coefficient^14^ to characterize confined diffusion of receptors. For comparison, HEK β-arrestin knockout cells transfected with cerulean-tagged β-arrestin2 were examined. In addition, HEK wild-type cells were analyzed in which β-arrestin2 recruitment was blocked by the GRK2 inhibitor paroxetine.

## Results and Discussion

### Validation of β_2_-adrenergic receptor labeling and cell model functionality

Wide-field fluorescent imaging of BG-CF640R labeled HEK SNAP-β_2_ cells was performed in wild-type and β-arrestin-KO genetic backgrounds (Fig. 1). Fluorescence intensity corresponding to stained SNAP-β_2_ adrenergic receptors was high along the plasma membrane of the cells. Fluorescent artefacts by non-specific binding were low and cell morphology normal, with no ostensible differences between both genetic backgrounds.

**Figure 1:**
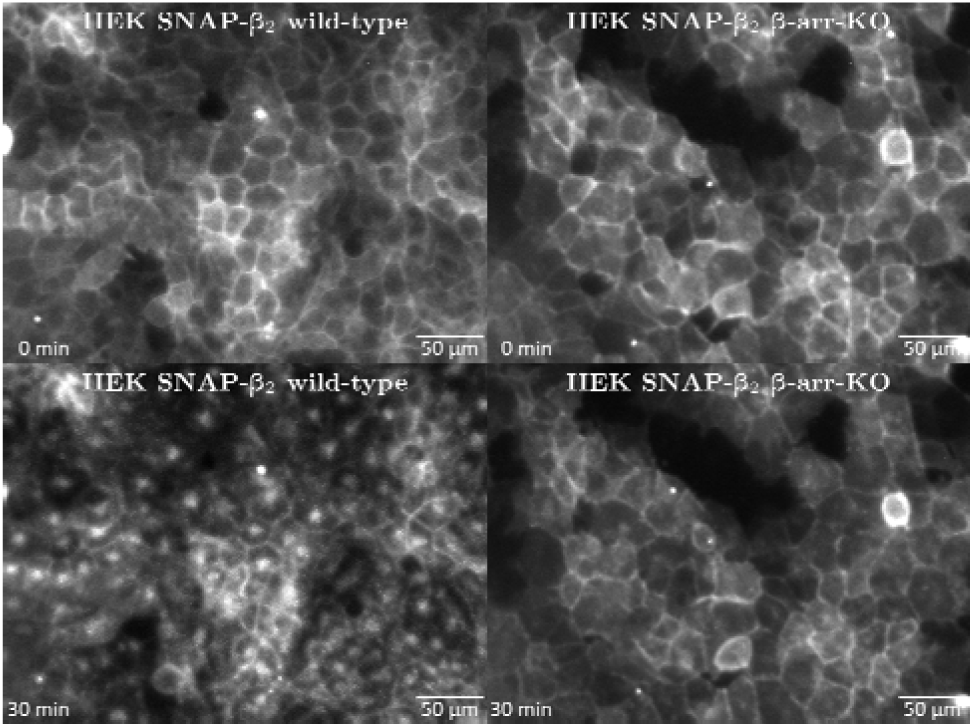
Fluorescent staining of HEK SNAP-β_2_ cells with BG-CF640R in a basal and stimulated state. Isoprenaline (10 μM) was added after the initial image acquisition at t = 0 min. **Top:** In the basal condition, plasma membranes were clearly stained in both genetic backgrounds, indicating expression and membranous location of β_2_AR. **Bottom:** Following stimulation in HEK SNAP-β_2_ wild-type cells, the membrane staining was displaced by endosomal receptor internalization, as seen in the formation of concentrated fluorescent spots within the cells. This effect was absent in the HEK SNAP-β_2_ β-arr-KO cells, which did not show differing receptor distribution following stimulation.

Stimulation of stained HEK SNAP-β_2_ wild-type cells with 10 µM isoprenaline, a β-adrenoceptor agonist, led to the formation of concentrated fluorescent spots within cells after 30 minutes, corresponding to early endosomes carrying labeled β_2_AR. Following stimulation in HEK SNAP-β_2_ β-arr-KO cells, fluorescence was still localized in the plasma membrane after 30 minutes, with no visible formation of early endosomes.

Both dye specificity to SNAP-β_2_AR and strongly visible stimulation response of the cells prove the suitability of this model system for single particle tracking. Further, it allows the introduction of newer and more precise evaluation methods, which will be done in the following.

### Reduced mobility of β_2_-adrenergic receptors after agonist stimulation

For single particle microscopy, the cells were labeled at a much lower dye concentration (10 nM) and incubation time (5 min) to achieve optical separation of single receptor molecules. The movement of single particles in the optically focused apical membrane of individual cells was clearly visible and was recorded for 50 seconds per cell at 20 Hz using an EMCCD camera. Diffusion behaviour of labeled receptors under non-stimulating conditions was recorded for 30 minutes. Subsequently, the cells were stimulated with isoprenaline in order to detect alterations in lateral mobility under stimulating conditions, again in a time frame of 30 minutes. Particles were later automatically localized and tracked (Fig. 2). The data of 160 cell measurements were obtained from eight independent experiments and were pooled in respect to genetic background and stimulation condition.

**Figure 2:**
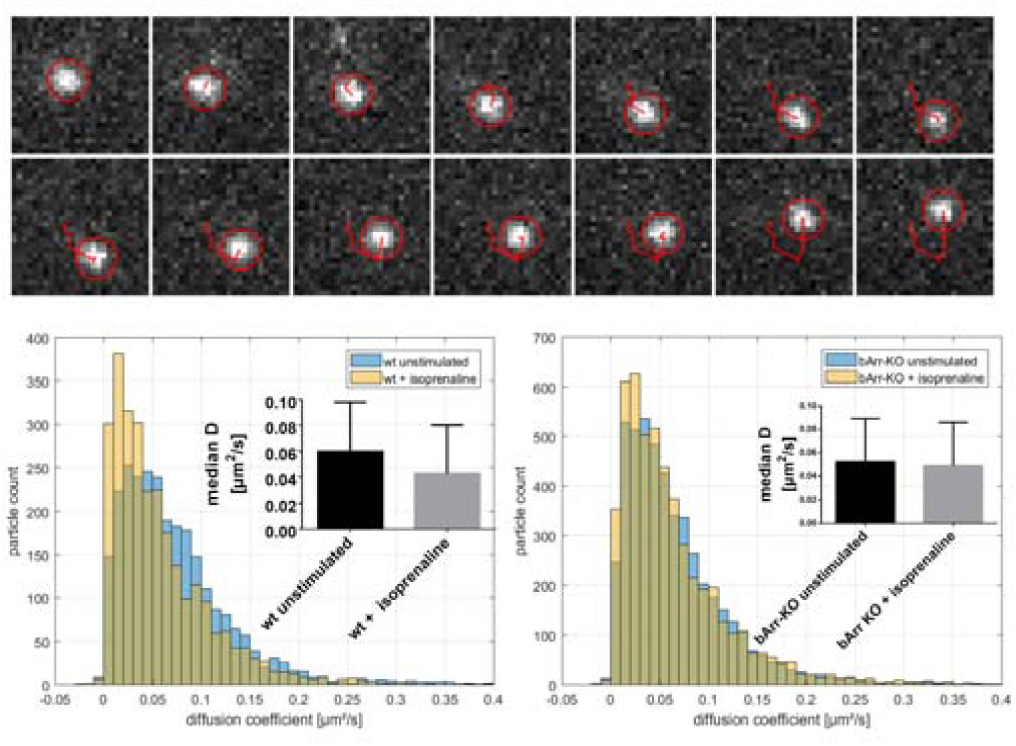
**TOP:** Example of particle localization and track creation. Shown is a cropped region of 2.6 × 2.6 μm in the apical cell membrane, recorded in 50 ms intervals, in which a freestanding single particle is diffusing. The fluorescent signal of this particle is detected and its origin precisely localized by Gaussian fitting. The coordinates are subsequently linked to resemble the path of lateral receptor movement. **Bottom left:** β_2_AR in HEK SNAP-β_2_ wild-type cells showed a heterogeneous diffusion coefficient distribution with a median D of 0.060 μm^2^/s. Following stimulation, the proportion of small (D < 0.4 μm^2^/s) diffusion coefficients strongly increased, reducing the median D to 0.043 μm^2^/s. **Bottom right**: β_2_AR in HEK SNAP-β_2_ β-arr-KO cells showed a similar distribution pattern with slightly lower basal (unstimulated) values, as indicated by a median D of 0.053 μm^2^/s. Stimulation resulted in only a slight shift towards lower diffusion coefficients, with a median D of 0.049 μm^2^/s.

The movement of particles was first analyzed by linear fitting of mean squared displacement curves to receive short-term diffusion coefficients of single particles (Fig. 2). In HEK SNAP-β_2_ wild-type cells the labeled β_2_AR showed a heterogeneous diffusion coefficient distribution with a median D of 0.060 µm^2^/s (N = 2700 tracks). Following stimulation, diffusion coefficients strongly shifted towards low values, reducing significantly the median D to 0.043 µm^2^/s (N = 2766). This effect likely corresponds to desensitized receptors prior to internalization (compare Fig. 1).

Evaluation of HEK SNAP-β_2_ β-arr-KO cells showed a comparable distribution of β_2_AR diffusion coefficients with a lowered median D of 0.053 µm^2^/s in the basal state (N = 4909). Under stimulating conditions only a slight shift toward slower diffusion coefficients was observed, with a median D of 0.049 µm^2^/s (N = 5147). Thus, the isoprenaline mediated reduction in median D of β_2_AR depends on the presence of β-arrestins, leading to receptor internalization together with other factors. To study the oligomerization states of β_2_AR detected on the cell surface, histogram analysis of the fluorescence intensities of all the particles were performed for each condition (Fig. 1, supplemental material). Data of HEK SNAP-β_2_ wild-type cells were fitted with a mixed Gaussian model. Four different intensity populations were observed corresponding to 31% of receptor monomers, 42% of receptor dimers, 21% of receptor trimers, and 6% of receptor tetramers. This finding was also found for HEK SNAP-β_2_ β-arr-KO cells and did not change even after isoprenaline stimulation for both genetic backgrounds (Fig 1, supplemental material). Thus, reduction in median D of β_2_AR under stimulating conditions cannot be attributed to increased formation of dimers. Calebiro et al. analysed β_2_AR oligomerization and lateral mobility using single-molecule TIRF microscopy^15^. They found for overexpressed SNAP-tagged β_2_AR in CHO cells fractions of mono-, di-, tri-, and tetramers with respective abundances of approx. 5%, 50%, 30%, and 15%. Remarkably, the distribution of the oligomerization states in CHO cells is not exactly the same as in our HEK cells. In particular, the presence of monomers was more pronounced in our experiments. Similar to our results Calebiro et al. did not find an effect on the oligomerization state of β_2_ARs after agonistic stimulation^15^.

The difference in particle numbers between both genetic backgrounds can most likely be explained by differing expression levels of β_2_AR, rather than β-arrestin pathway blockage. Hence, direct comparison of both backgrounds is more ambiguous than analyzing the effects of stimulation, which is prioritized in our experiments.

In a similar experiment that used SNAP-tag labeling of β_2_AR in CHO cells, a comparable median D of 0.039 µm^2^/s was determined by Calebiro and co-workers^15^. Remarkably, isoprenaline stimulation using the same concentration had no effect on diffusivity, which was not further discussed by the authors. Sungkaworn et al. also did not observe significant changes to diffusion states in similar conditions^4^. We think that in our experiment, endogenous expression of β_2_AR in HEK^16^ or high levels of receptor expression in stable clones may be the deciding factors that lead to receptor desensitization and altered diffusion. Our results agree with a recent study by Yanagawa et al., in which the average MSD-derived diffusion coefficient of β_2_AR in HEK cells was reduced from 0.078 to 0.051 µm^2^/s by stimulation using 10 µM isoprenaline^17^. This suggests that principally changes in diffusion coefficient are comparable even for different imaging modalities (TIRFM and wide-field) and receptor expression strategies (transient expression and stable clones).

### A three-state classification model showed consistent values for diffusion coefficients obtained by vbSPT evaluation

The diffusion of β_2_AR was further analyzed by applying an algorithm based on variational Bayesian treatment of a hidden Markov model (vbSPT). In contrast to previous hidden Markov model approaches for single molecule analysis, vbSPT is capable to extract useful information even from data sets with short trajectories. vbSPT also allows the detection of state transition probabilities, as well as the number of diffusive states from experimental data. Trajectories were segmented as described by Persson et al. and segments were classified to one of three distinct diffusion states (S1 to S3)^18^. Each of these states is defined by diffusion coefficient, occupancy value and state switching probabilities. To maximize model accuracy, data were pooled by condition as before, with 40 cells per analysis. The resulting state diagrams (Fig. 3) showed a similar model in all four conditions. Diffusion coefficients for each state were nearly constant in all conditions and ranged from 0.009 to 0.011 µm^2^/s (S1), 0.035 to 0.038 µm^2^/s (S2) and 0.116 to 0.123 µm^2^/s (S3). This consistency demonstrated model robustness and allowed comparison of other metrics associated with the model, namely state occupancy and transition probabilities. For bar graphs and significance testing (Figure 3), slightly altered data was produced by batch wise data splitting and analysis (4 × 10 cells per condition).

**Figure 3:**
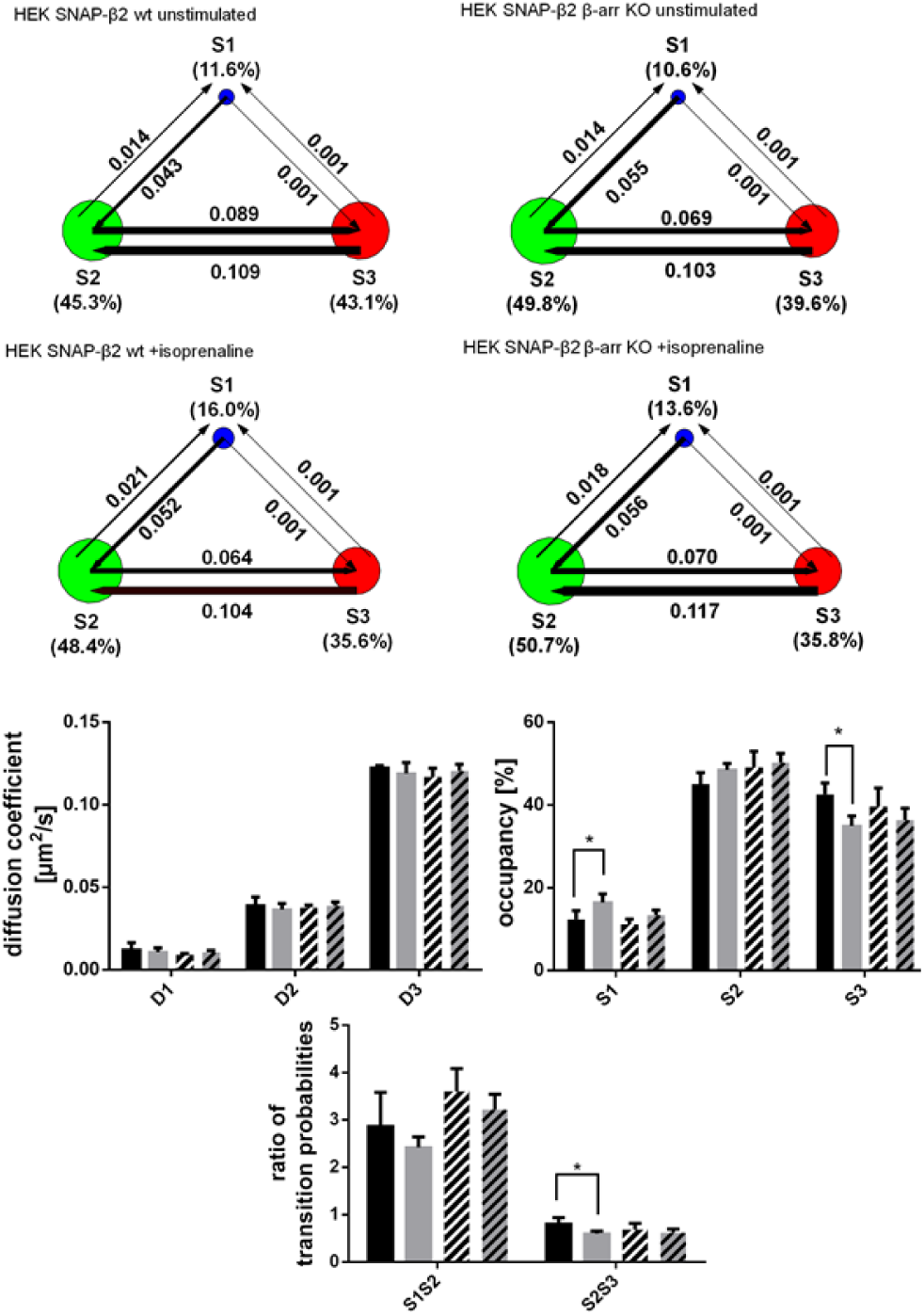
Diffusion state models for all conditions, attained by vbSPT analysis of pooled track data with HEK SNAP-β_2_ wild-type cells before (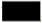, 13126 tracks) and after (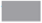, 14429 tracks) stimulation and HEK SNAP-β_2_ β-arr-KO cells before (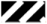, 19198 tracks) and after (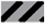, 21961 tracks) stimulation. Coloured circles represent discrete diffusion states S1 to S3 and circle size indicates occupancy. Arrows indicate switching rates between states. Marked value with p ≤ 0.05 is significantly different from its corresponding control, determined by Student’s t-test.

### Agonist stimulation altered receptor state occupancies by state switching

In HEK SNAP-β_2_ wild-type cells that were stimulated with isoprenaline, occupancy of slow-diffusing state S1 increased from 11.6 % to 16.0 %. Similarly, the fraction of receptors in S2 increased from 45.3 % to 48.4 %, whereas the fraction of receptors in fast state S3 was strongly reduced from 43.1 % to 35.6 %. Only the changes for S1 and S3 occupancy were significant (Fig. 3).

The same trends were seen in HEK SNAP-β_2_ β-arr-KO cells, but to a lesser extent: Stimulation increased S1 occupancy from 10.6 % to 13.6 %. Occupancy of S2 was initially higher at 49.8 %, and only increased to 50.7 %. The occupancy of S3 was reduced from 39.6 % to 35.8 %.

Although stimulation initiated the same trends that were seen in HEK SNAP-β_2_ wild-type cells, the changes here were not significant. Obviously, the subsequent regulation of activated β_2_AR in HEK SNAP-β_2_ β-arr-KO cells was disturbed, as seen in the missing internalization of β_2_AR under stimulating conditions (Fig. 1).

By transfection of cerulean-tagged β-arrestin2 into HEK SNAP-β_2_ β-arr-KO cells, we were able to demonstrate that the observed effects depend on the absence of β-arrestin. HEK SNAP-β_2_ β-arr-KO cells expressing β-arrestin2 were identified by the cerulean-tag in the fluorescence microscope. After isoprenaline stimulation of these cells (n=10) in single particle tracking experiments, the occupancy of S1 increased significantly from 15.7 % to 23.6 %, whereas S2 and S3 were slightly reduced from 54.9 % to 51.1 % and 29.4 % to 25.2 %, respectively (Fig. 4). Furthermore, the number of trajectories decreased from 8577 to 5214, suggesting an internalization of β_2_AR. Accordingly, transfected HEK SNAP-β_2_ β-arr-KO cells expressing cerulean-tagged β-arrestin2 showed HEK SNAP-β_2_ wild-type cell like behaviour.

**Figure 4:**
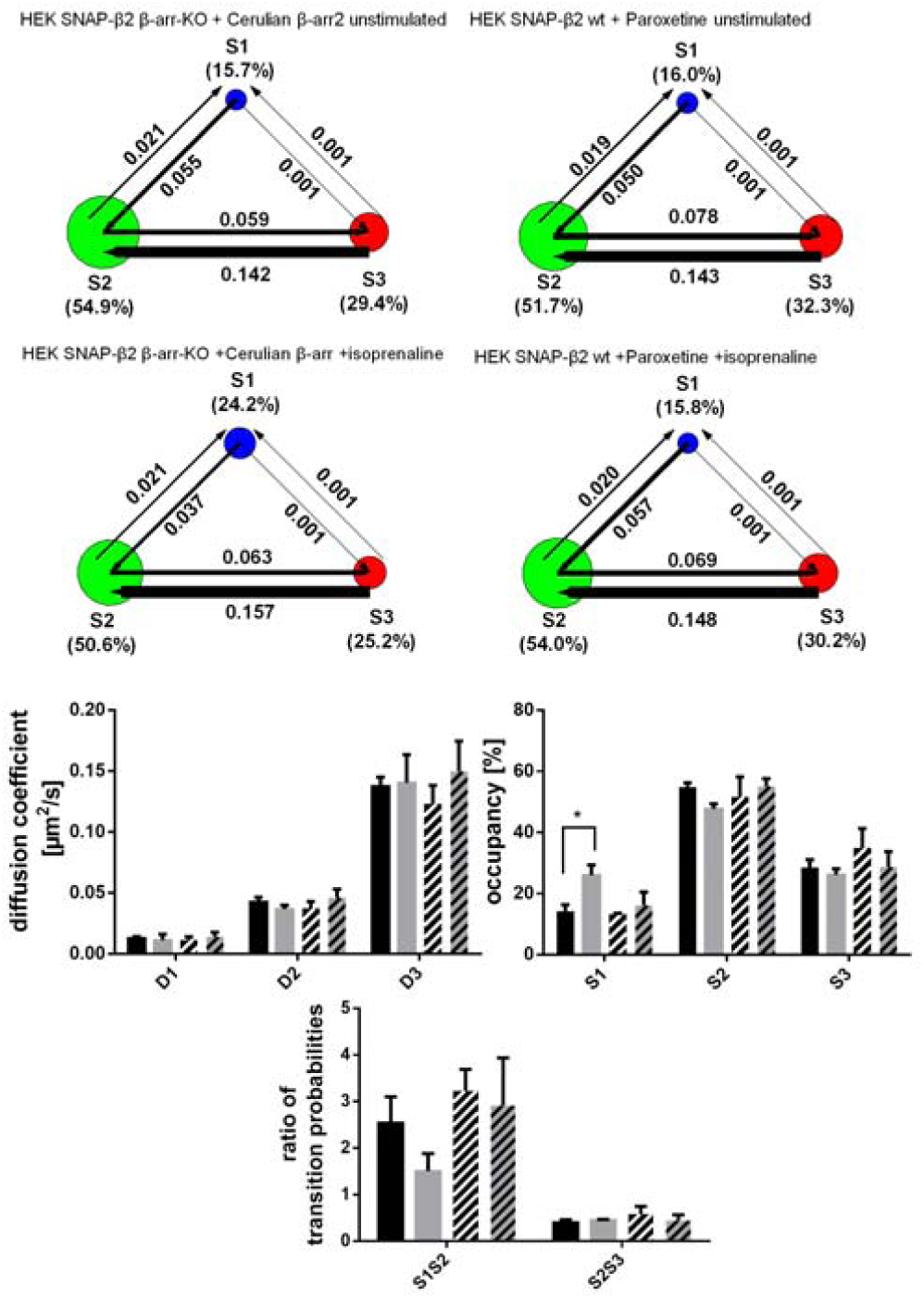
Diffusion state models for all conditions, attained by vbSPT analysis of pooled track data with HEK SNAP-β_2_ β-arr-KO cells transfected with cerulean-tagged β-arrestin2 before (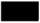, 8577 tracks) and after (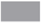, 5214 tracks) stimulation and HEK SNAP-β_2_ wild-type cells pre-incubated with paroxetine before (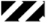, 5636 tracks) and after (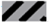, 5263 tracks) stimulation. Coloured circles represent discrete diffusion states S1 to S3 and circle size indicates occupancy. Arrows indicate switching rates between states. Marked value with p ≤ 0.05 is significantly different from its corresponding control, determined by Student’s t-test.

We propose that slowed diffusion by altered state occupancies of activated β_2_AR (Fig. 3, 4) is caused by the molecular interactions involving β-arrestin, further supported by lack of a significant stimulation response in HEK SNAP-β_2_ β-arr-KO cells. The importance of β-arrestin for the signalling of β_2_AR was demonstrated by influences of paroxetine, a GRK2 inhibitor, on β_2_-adrenergic signal transduction. Guo et al. observed that paroxetine inhibits the isoprenaline-induced phosphorylation of β_2_AR, the recruitment of β-arrestin2, and the subsequent receptor internalization^19^. In our experiments paroxetine pre-treated and isoprenaline stimulated HEK SNAP-β_2_ wild-type cells changed the occupancy of S1 from 16.0 % to 15.8 %, S2 from 51.7 % to 54.0 % and S3 from 32.3 % to 30.2%, which was not significant (Figure 4). Since the trajectory number after isoprenaline stimulation decreased only slightly from 5636 to 5263, paroxetine also inhibited the internalization of β_2_AR in these cells as described above. Thus, paroxetine pre-treated HEK SNAP-β_2_ wild-type cells behaved like HEK SNAP-β_2_ β-arr-KO cells.

The observed differences in occupancy between conditions correlate with altered probabilities of state switching of the registered particles (shown by arrows in Fig. 3 and 4). Two general observations were made: Regardless of genetic background, pre-treatment, and stimulation, state switching rates between S1 and S3 were neglectable (P_13_ and P_31_ <= 0.001), thus proving sequentiality of state switching. Secondly, P_23_ and P_32_ were consistently much higher than P_12_ and P_21_, indicating that receptors were more likely to change between the two fast states.

Transition probability ratios between states were calculated as the ratio of forward to backward switching probabilities between adjacent states and revealed significant changes for S2↔S3 transitioning in HEK SNAP-β_2_ wild-type cells following stimulation (Figure 3). A similar finding was observed by Ibach et al. for SNAP-tagged epidermal growth factor receptor in MCF-7 cells^20^. Since vbSPT was developed based on processes in thermodynamic equilibrium, we agree with the authors that transition probability data have to be interpreted with care. The reduced ratio indicated a stronger preference of receptors for S2 compared to S3. This change was also present, but not significant under β-arrestin-KO. The ratio of S1↔S2 transitioning was lowered in both genetic conditions under stimulation, but the change was not significant. HEK SNAP-β_2_ β-arr-KO expressing cerulean-tagged β-arrestin2 showed a wild type similar decrease in the ratio of S1↔S2 transitioning, but no difference in the ratio of S2↔S3 transitioning.

### Diffusion states differed in spatial confinement

To further characterize the nature of each diffusion state, the track segments corresponding to each state were extracted from the full trajectories and subjected to a confinement analysis based on their state and packing coefficient (Pc). The Pc values were calculated for sufficiently long segments using a 10 frame (0.5 s) sliding window, and compared against a 5 % false detection likelihood threshold (Pc_95_) based on the simulation of random diffusion.

The results in HEK SNAP-β_2_ wild-type cells are shown in Figure 5. In basal condition, about two-thirds (68.2 %) of track segments of S1 showed a confined diffusion with a mean confinement size (square root of convex hull area) of 59 nm. Particles in S2 were less often confined (42.7 %) and adhered to a larger confinement size of 102 nm on average. The fast state S3 showed an even smaller fraction of confined trajectory segments (28.2 %) with an increased confinement size of 182 nm.

**Figure 5:**
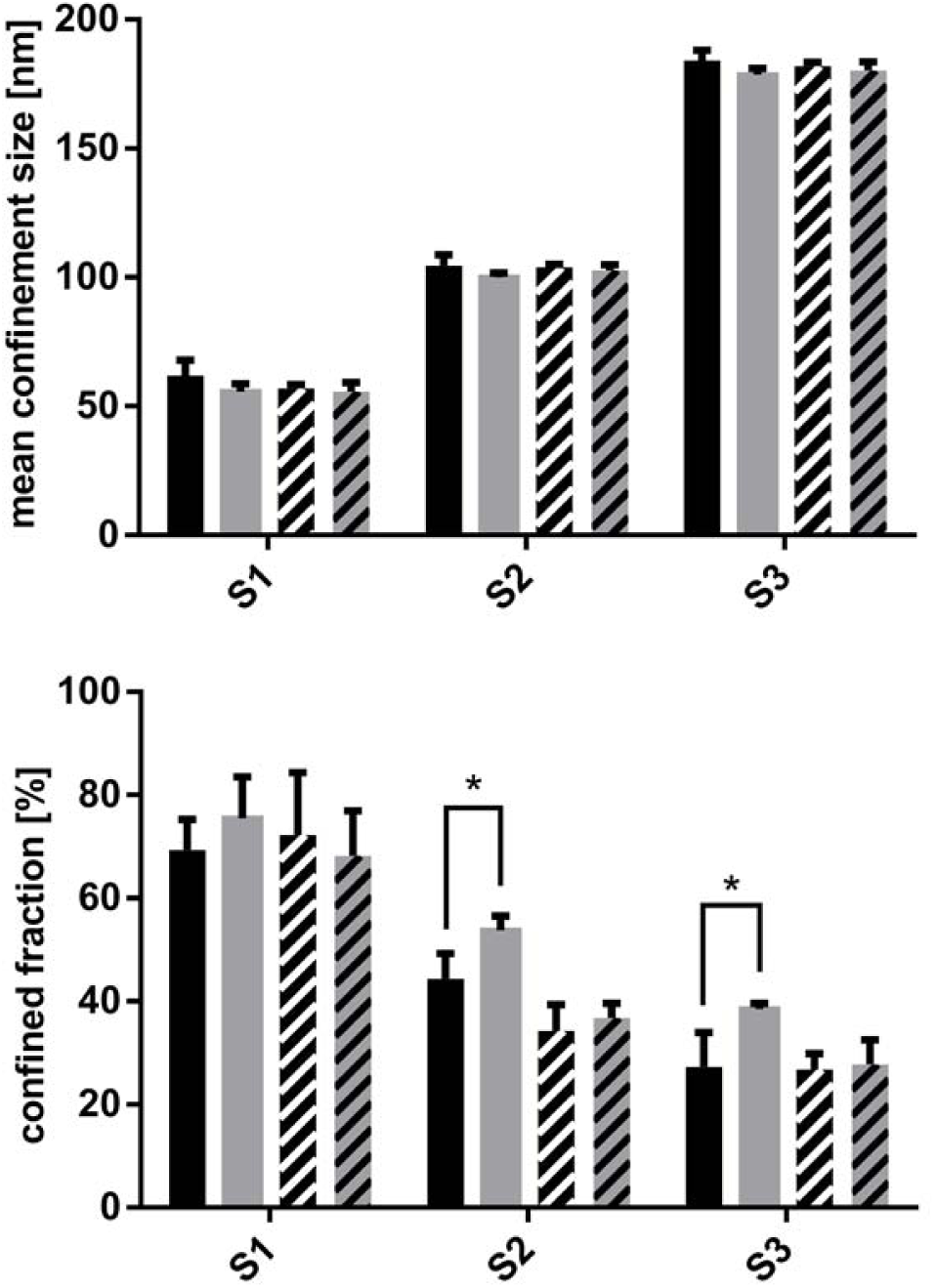
Mean confinement size and confined fraction of HEK SNAP-β_2_ wild-type cells before (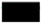) and after stimulation (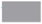) and HEK SNAP-β_2_ β-arr-KO cells before (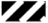) and after stimulation (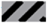) attained by Pc analysis. Marked value with p ≤ 0.05 is significantly different from its corresponding control, determined by Student’s t-test.

The fact that all states included a degree of confined diffusion and that mean confinement areas were nearly constant for each state regardless of genetic background and stimulatory condition are noteworthy, as previous research often classified states rigidly as either confined or free diffusion.

Unlike the high stability of state diffusion coefficients and also confinement areas, which were hardly changed by isoprenaline stimulation, the confined fraction was increased in each state. Confined diffusion was expected based on the recruitment of β_2_AR for the internalization process under stimulating conditions (Fig. 5). This process however does not correspond to the confined fraction of one specific receptor state, since all confined fractions are affected by stimulation. Knockout of β-arrestin (Figure 5) lessened the increase of confined fractions following stimulation and even slightly reduced it in S1. Further experiments will hopefully help to establish more detailed biological correspondence for each state. For example, these experiments may use shorter time periods following stimulation, monitor the molecular interaction of receptors, or analyze the diffusion of bound ligands.

## Conclusion

The HEK SNAP-β_2_ cell system with BG-CF640R labeling was well suited to investigate localization, lateral membrane diffusion and associated confinement patterns of β_2_-adrenergic receptors. In addition to a general characterization, stimulation with 10 µM isoprenaline evoked strong changes that could be tracked in a macro- and micromolecular scale by using increasingly sensitive methods: In short, classical fluorescence microscopy showed internalization of membraneous β_2_AR following stimulation. Histogram analysis of the fluorescence intensities obtained from SPT of β_2_AR showed receptor mono-, di, tri, and tetramers on the cell surface. In MSD-based analysis of single particle tracks, receptor internalization was reflected by slowed diffusion in a heterogeneous diffusion coefficient distribution. MSD is a classical approach that is heavily averaging and therefore does not reflect all information that is contained in particle trajectories. Since the plasma membrane is a complex and heterogeneous environment that continuously influences the diffusion of β_2_AR, there is a need for a more sophisticated analysis to resolve the complex lateral mobility.

By variational Bayes analysis, we found three underlying diffusion states for β_2_AR: A slow state, an intermediate state and a fast state, with constant confinement sizes (56 nm, 105 nm, 184 nm) in each state even under stimulation. Surprisingly, all states including the fast diffusive state had a confined fraction of receptors. These fractions were changed following stimulation, as well as state occupancies, a change that was not seen in previous research.

The changes to occupancies result from altered state switching that is exclusively sequential via intermediate state S2. Inter-state transitioning probabilities were strongly in favor of S2, also underlined by the observation of low occupancy in the slow state S1 and high affinity for switching to S2.

The data and methods presented contribute to a more precise characterization of physiological states and molecular interactions of β_2_AR. We believe that our findings enable the research of fine-grained differences between other conditions that manipulate β_2_AR signaling and can easily be adapted to other GPCRs.

## Material and Methods

### Dye Preparation

The fluorescent SNAP-tag substrate BG-CF640R was synthesized by amidization reaction of BG-NH_2_ (New England Biolabs #S9148S) and CF640R-NHS ester (Biotium #92108). High-performance liquid chromatography was conducted to purify the desired reaction product (see Supporting Information), as verified by subsequent MALDI-TOF measurement (m/z = 1085.24).

To enrich BG-CF640R, the corresponding HPLC fraction was subjected to a RP-18 solid phase extraction. Phosphoric acid-containing eluent was removed by washing with water. BG-CF640R was then eluted with a mixture of methanol and ethanol. The eluate was dried using a vacuum centrifuge.

A stock solution of BG-CF640R in DMSO was prepared to a concentration of 400 µM and stored at - 20°C. For labeling, the stock solution was diluted in water to a concentration of 4 µM and stored at 4°C.

### Cell culture and transfections

Human embryonic kidney (HEK) cells were obtained from DSMZ (Braunschweig, Germany). HEK β-arrestin-KO cells were kindly provided by Asuka Inoue (Graduate School of Pharmaceutical Science, Tohoku University, Sendai 980-8578, Japan). Both cell lines were maintained at 37°C and 5% CO_2_ in DMEM medium (Gibco #31885-023) containing 10% fetal calf serum (Life Technologies #10270), 100 units/ml penicillin and 100 µg/ml streptomycin.

The plasmid coding for SNAP-β_2_AR was obtained from New England Biolabs (#N9184). Transfection was done by calcium phosphate transfection method: Cells were seeded in 12-well plates and allowed to attach for at least 24 hours. One µg plasmid DNA was mixed with 6.5 µl of a 2M aqueous CaCl_2_ solution and 50 µl sterile water, and then added dropwise to a two fold HBS buffer (pH 7.13, 42 mM HEPES, 274 mM NaCl, 10 mM KCl, 1.4 mM Na_2_HPO_4_, 15 mM glucose). The mixture was added to the cells after 30 minutes. On the next day, the medium was changed to fresh DMEM containing 750 µg/ml G418 for selection. Individual clones were selected in cloning rings and seeded in distinct wells in a 12-well plate. The clone with the best expression of SNAP-β_2_AR was identified by fluorescence microscopy and used for all experiments.

To obtain β-arrestin2 expression vectors with N- and C-terminal fusion to Cerulean, respectively, sequence information for β-arrestin2 was amplified by PCR using the plasmid pECFP-N1_rβ-Arrestin-2 (a kind gift from Prof. Lohse – University of Wuerzburg, Germany^21^) as a template. PCR products were cloned into pCerulean-N and pCerulean-C by using NheI and AgeI restriction sites. All constructs were verified by DNA sequencing. DNA was transiently transfected into HEK SNAP-β_2_ β-arr-KO cells by electroporation with the Amaxa Nucleofector II device (Lonza). The included program Q-001 is designed for high expression levels in HEK cells and was used according to the manufacture’s protocol. Single particle microscopy experiments were carried out three days after transfection. Cells expressing cerulean-tagged β-arrestin2 were identified by fluorescence microscopy and used for all experiments.

### Fluorescence staining and microscopy

For experiments, cells were seeded in 12-well plates on fibronectin-coated glass coverslips using clear DMEM medium (no phenol red, Gibco #11880-028). Coating was performed by pre-incubating wells prior to seeding, using 500 ng/µl fibronectin in PBS for two hours at 37°C, then washing twice with PBS. Fibronectin coating improved cell adhesion and growth compared to poly-D-lysine coating, especially in the knockout cell line.

Experiments were performed two or three days after seeding, at a confluency of about 80%. For fluorescence microscopy, SNAP-β_2_AR over-expressed in HEK cells were fluorescently labeled by preparing a solution of 2.5 µM BG-CF640R in clear medium and incubating at 37°C and 5% CO_2_ for 30 minutes. This was followed by three washing steps with clear medium, and a change to HBSS buffer (Gibco #14025050) for imaging. The coverslip was then placed in a custom made mounting bracket and imaged at 25°C.

### Single particle microscopy

Single particle microscopy was performed using an inverted wide-field epi-fluorescence microscope (TE2000-S, Nikon, Kanagawa, Japan), equipped with a 60x water-immersion objective (Plan Apo VC, 1.2 NA, Nikon), a 200-mm-focal-length tube lens, a 4x-magnification lens (VM Lens C-4x, Nikon) and a EMCCD camera (iXon DV-860DCSBV, Andor Technology). For fluorescence excitation, a 637 nm continuous wave laser (Coherent) was set to 20% intensity 0.7 kW/cm^2^ in the object plane) using an acousto-optical tunable filter (A.A SA, France). The recording conditions were kept constant for all measurements (1000 frames at 20 Hz, continuous illumination and camera exposure, constant camera gains and readout speeds).

The SNAP-tag labeling procedure described above was altered to achieve sparse labeling. For appropriate spot densities and lower background signal, a five minute incubation using only 10 nM SNAP-tag dye was optimal. After labeling and washing, the cells were immediately imaged at 20°C and used no longer than 60 minutes. In this time, 20 individual cells were recorded using the following procedure: A cell was centrally positioned in the bright-field channel and focused to the apical membrane. Then, the EM channel and laser were activated and the focus briefly fine-tuned before starting the image acquisition. Cells showing either unusually low spot densities or areal fluorescent artifacts were generally discarded.

### Spot tracking

The MATLAB software (version R2016b, MathWorks) was used for the generation of 2D particle tracks from image data and further diffusion analysis. Images were directly imported by the u-track package^22^ and processed using the following settings: 1.32 px spot radius, 3 frame rolling window time-averaging for local maxima detection, 2 frame minimum track segment length, 1 frame maximum gap length, other settings on default.

Gaps were afterwards closed by linear interpolation, and the resulting particle tracks pooled by experimental condition and evaluated using scripts described in the following.

### Analysis of diffusion behavior

We implemented the package *@msdanalyzer*^23^ to calculate mean squared displacement (MSD) values for each track. MSD is calculated as in eq. 1, with r(t) as the particle’s position at time t, N as number of positions in the trajectory and *m*Δ*t* as the variable time lag, a multiple of the frame interval.

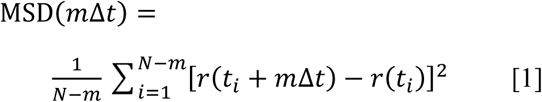

Short-term diffusion coefficients of labeled receptors were then derived by linear fitting of the first four MSD points (eq. 2), weighted by number of distances in each MSD value. Tracks of less than 8 frames or low fit goodness (r^2^_adj_ < 0.85) were excluded.

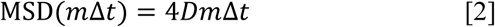

To identify discrete diffusive states from particle tracks, variational Bayes single particle tracking was applied using the vbSPT Matlab package^18^. Tracks were thereby segmented and variably classified to one of three states according to their momentary diffusion speed. Higher order models were recognized by the program but not used, since they resulted in degenerate states of insignificant occupancy and indistinct diffusion behaviour (Figure 2, supplemental material).

The introduction of a fourth state to the model leads to the addition of a nearly unpopulated immobile state S1, and the detection of higher diffusion coefficients in the other three states. MSD analysis of the segments corresponding to each state showed that on longer timescales, S1 and S2 in the four-state model were degenerate, since short- and long-term diffusion were extremely similar. Higher order models also showed one or more degenerate states.

### Track simulations and confinement analysis

Tracking data resembling Brownian molecular motion was simulated as follows: 500.000 tracks were generated for each diffusion coefficient (D_1_ = 0.011 µm^2^/s, D2 = 0.038 µm^2^/s, D3 = 0.12 µm^2^/s) at a frame interval of 50 ms. All generated spots were subjected to localization error by a normally distributed positional offset with σ = 20 nm in each dimension. To account for photobleaching, the track lengths were modeled by an exponential distribution with μ_D1_ = 21.3 frames, μ_D2_ = 9.8 frames, μ_D3_ = 7.1 frames with a minimal track length of 2 frames. Values for D and µ were derived from averaged vbSPT analysis results of the real data (β_2_AR under non-stimulating and stimulating conditions).

For the analysis of confinement, the previously classified track segments were extracted and pooled by their respective diffusion states. The recently introduced packing coefficient Pc^14^ was used as a measurement of spatial confinement strength. It is defined in a given time window as the sum of squared displacements divided by the squared convex hull area formed by the included particle positions. A window length of 10 points (0.5 s) was chosen, which should be long enough to yield stable results (see Fig. 5 in ^14^) and still include sufficient numbers of track segments. To determine Pc_95_-values given by the 95th percentile of packing coefficients, random walk data based on the previously determined vbSPT state diffusion coefficients and segment lengths was simulated. The derived Pc_95_-values were then used as a threshold for spatial confinement. Confined tracks (Pc > Pc_95_) were then compared by their average hull areas, again by averaging on the 0.5 s timescale.

## Abbreviations

(SPT): single particle tracking
(β-arr): β-arrestin
(β_2_AR): β_2_-adrenergic receptor

## Acknowledgments

We thank Asuka Inoue (Graduate School of Pharmaceutical Science, Tohoku University, Sendai 980-8578, Japan) for kindly providing HEK β-arrestin-KO cells.

**The authors do not declare any conflict of interest**

## Notes

#### Summary of Updates

The manuscript was reviewed and the obtained comments led to a substanstial improvement of the manuscript.

