## Supplemental Material for "Live-cell single-molecule analysis of β_2_-adrenergic receptor diffusion dynamics and confinement"

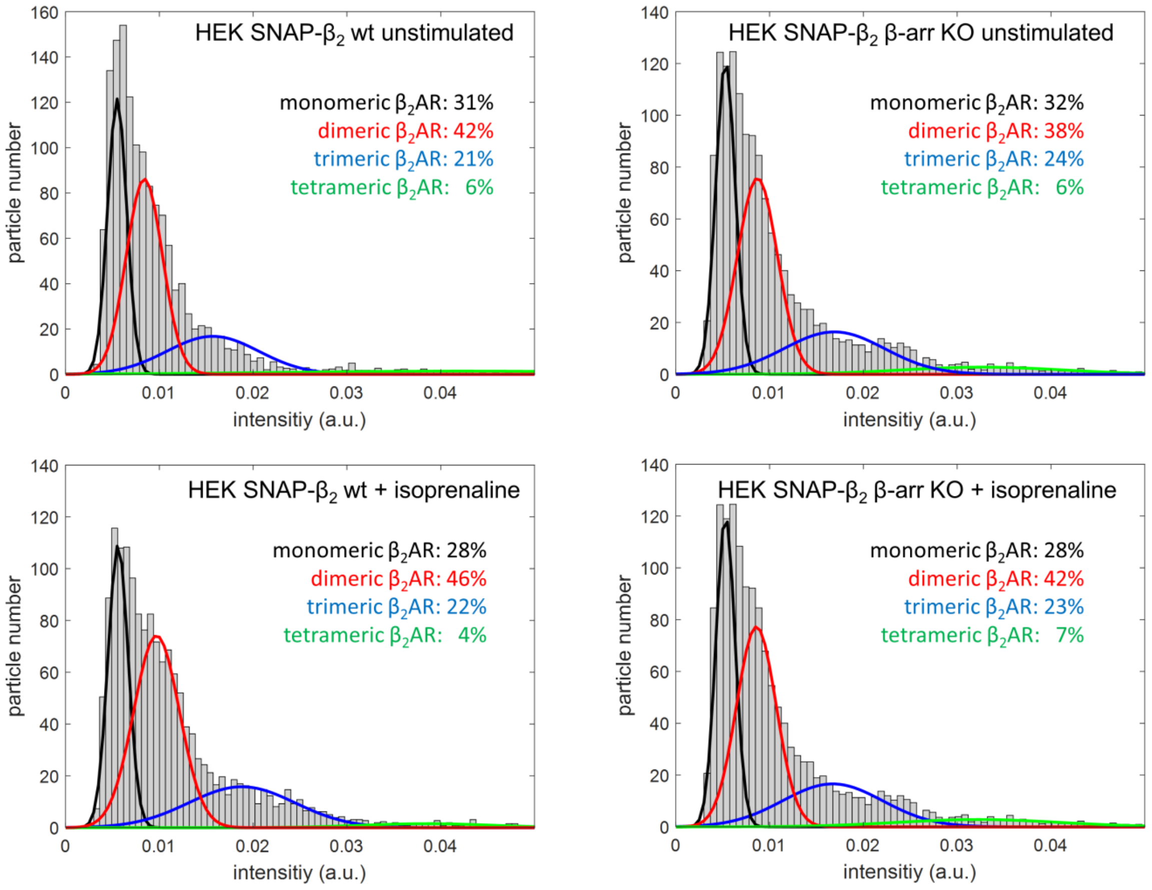


**Figure 1:** Histogram analysis of fluorescence intensities of SNAP-tagged $\beta_2$AR analyzed in each condition. $\beta_2$AR oligomerization states were obtained from data fitting using a mixed Gaussian model.

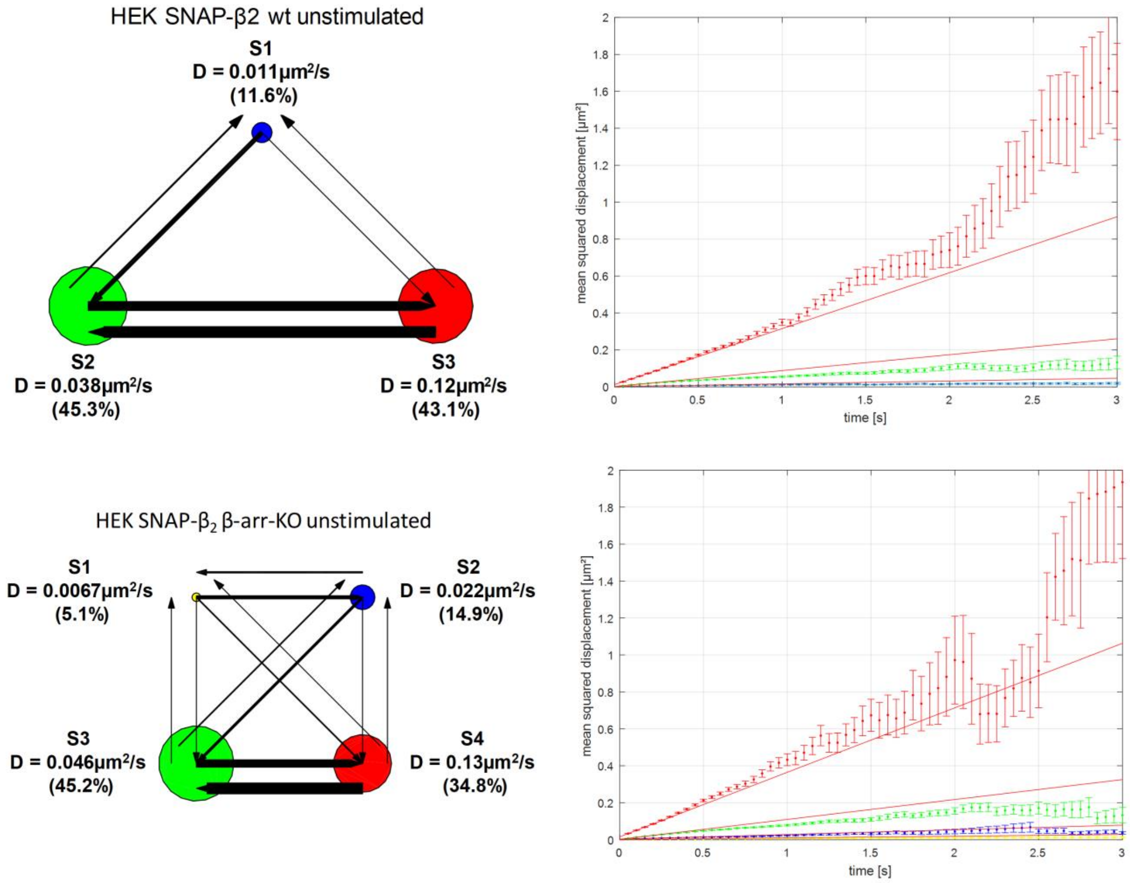


**Figure 2**: Left: Comparison of a three-state model (top) and four state model (bottom) generated by vbSPT analysis of non-stimulated HEK SNAP-$\beta_2$ wild-type cells. State diagrams are shown in which circle sizes indicate occupancy. Arrows indicate switching rates between states. Right: MSD analysis of the segments corresponding to each state for both the three- and four state model.
